# What’s in a Story? Studying Neural Narrative Engagement with Auditory Passages Using EEG Inter-Subject Correlation

**DOI:** 10.64898/2026.11.19.767982

**Authors:** Madison Bunderson, Blair Kaneshiro, Philip A. Hernandez, Neha Rajagopalan, Bruce D. McCandliss

## Abstract

Narrative engagement — a state implicating attention as well as emotional and sequential processing — is crucial in contexts ranging from social interaction to education. Yet we still lack an understanding of how narrative contents and listener perceptions relate to the neural responses they elicit. In this study, we investigate relationships between auditory narratives and listener responses using electroencephalography inter-subject correlation (EEG-ISC), a measure of neural synchrony that has been shown to correlate with narrative cohesion, attention, preference, and learning. We analyzed dense-array EEG recorded from 46 adult participants who listened to social and non-social narrated literary excerpts. The data were spatially optimized using Reliable Components Analysis, and ISC was calculated for each of the three maximally reliable components (RCs) on a per-stimulus basis. While we found no significant relationships between EEG-ISC and broad social/non-social categorizations of the excerpts, ISC for high sociocognitive word usage significantly exceeded that of medium and low word usage in the third spatial component, RC3. We further observed a significant positive correlation between participants’ summed ratings of interest, compulsion, and enjoyment and ISC, both for RC2 and when summed across RC1–RC3. Interaction analyses suggest that passage content and behavioral responses contributed independently as additive effects to these relationships. Taken together, these findings extend previous research utilizing EEG-ISC in natural speech contexts and, importantly, suggest that the decomposition of EEG into maximally correlated spatial components can separate co-occurring neural processes engaged during story listening.

## 1 Introduction

> *“In the modern, information-saturated world, sustaining attention on a single continuous information stream, even in the absence of competing sensory stimuli, is a constant challenge”* (Ki et al., 2016).

Attentional engagement in our everyday lives is a complex, multifaceted state that varies in task demands depending on our current context (Ki et al., 2016). In education, learners are often expected to engage with auditory streams of information throughout their development; these auditory-based inputs may range from a read-aloud in the lower grades to an hour-long lecture in the college classroom. Understanding how individuals engage with narrative content is therefore crucial for both theoretical and applied content.

Narrative engagement, a specialized type of attentional engagement, refers to the psychological state of being absorbed in a story, and has been characterized in research by increased attention, emotional involvement with the characters and/or plot, and sociocognitive processing of the sequence of events (Busselle and Bilandzic, 2008). As every narrative progresses, there are multiple levels of narrative structure that can influence engagement — from moment-to-moment word level changes to overarching general plot structures. The brain processes these narrative structures hierarchically, accumulating information over long timescales through slow-frequency neural synchrony that integrates early story elements with later narrative developments (Honey et al., 2012; Lerner et al., 2011). Studies of literature have found that narrative features, ranging from the smallest details to larger, genre-specific conventions, can result in shifts in engagement (Busselle and Bilandzic, 2008; Cheng, 2008).

Different narrative features are ripe for further investigation, as they provide manipulable yet still naturalistic variables involving both genre (a broad categorization) and content (a more specific variable). In the current study, we chose to focus on two defining aspects of genre by examining the distinction between social and non-social texts. We identified this dimension because it is among the most frequent divisions used in the study of text and cognition (e.g., Black and Barnes, 2015; Colman, 2007), although these divisions often lack clear boundaries for categorizing texts. The division between social and non-social texts is driven by a plethora of information around the importance of social and sociocognitive features of literature in processing, which tend to feature more heavily in fiction versus non-fiction texts (e.g., Mar and Oatley, 2008). In the current study, we focus on sociocognitive vocabulary and characters and are especially interested in the impact of this division given ample evidence that social and non-social information utilize different underlying sociocognitive processes and neural networks (e.g., Tompson et al., 2019). Critically, during naturalistic listening to auditory narratives, Yeshurun et al. (2017) demonstrated that processing social information, especially as related to characters’ mental states and relationships, evoked highly synchronized neural responses across regions associated with social cognition. This suggests that social content in narratives drives shared patterns of neural engagement, making the social/non-social distinction particularly relevant for understanding how different types of narrative content modulate brain synchrony.

Foundational work to understand these sociocognitive processes began in functional magnetic resonance imaging (fMRI) research. By focusing on the distinction between social and non-social information, researchers have established that there is underlying neural network segregation between social cognition and spatial cognition (Saxe et al., 2004). More specifically, fMRI studies have shown that sentences rich in theory of mind (ToM) content (which involves social inference about a character) activate a distinct network related to reasoning about a character’s thoughts and beliefs (e.g., Dodell-Feder et al., 2011). During naturalistic narrative comprehension, these processes engage what is often referred to as the “theory of mind” network, a set of brain regions specialized for reasoning about others’ mental states (Saxe et al., 2004). This network, which partially overlaps with but is distinct from the broader default mode network, undergoes dynamic reconfiguration as listeners integrate character information and social context across the story’s timeline (Simony et al., 2016). These findings are based on carefully crafted and brief vignettes and sentences. This begets questions around more naturalistic stimuli — how might the brain interact with longer narratives across a period of sustained engagement?

Despite the frequency with which human listeners engage with highly dimensional and sustained auditory stimuli, engagement has typically been studied in laboratory settings (e.g., O’Connell et al., 2009) with stimuli that are much more artificial than those with which we engage outside the lab. As neuroimaging tools have evolved and the field of neuroscience has become more interested in naturalistic stimuli, approaches to studying how the brain processes and engages with narratives have become newly possible routes of inquiry. A seminal early study by Hasson et al. (2004) demonstrated that fMRI inter-subject correlation (ISC) could capture shared neural responses during movie viewing. This finding helped to establish a foundation for naturalistic neuroscience research. Subsequent ISC research using audio narratives demonstrated key findings about auditory story comprehension; Stephens et al. (2010) found that successful communication during auditory storytelling creates neural “coupling” between speaker and listener brains, with anticipatory synchrony (the prediction of upcoming words) serving as a primary driver of comprehension. Additionally, shared neural responses to auditory stories persist even when audio is partially masked by noise, suggesting that the brain prioritizes semantic meaning over auditory clarity when possible (Irsik et al., 2022).

Building on this foundation, electroencephalography (EEG) ISC has emerged as a complementary approach to studying narrative engagement. ISC is thought to index engagement because it measures the degree to which individuals’ neural responses are synchronized during stimulus processing. With this metric, higher synchronization suggests that the stimulus is driving similar sociocognitive and attentional processes across individuals. This synchronization is interpreted as shared engagement with the stimulus’s content (e.g., Dmochowski et al., 2012). Importantly, the relationship between neural synchrony and comprehension is content specific: listeners who interpret a story’s meaning similarly show higher brain-to-brain synchrony than those who interpret it differently (Nguyen et al., 2019), and this shared understanding reflects synchronized neural response rather than shared sensory input alone. In EEG-ISC research using audio-only narratives, Ki et al. (2016) found that audio-only stimuli with a clear narrative elicited reliable responses, and the modulation of the response was most strongly impacted by what the experimenters deemed an overall mood of the narrative (e.g., suspense; Ki et al., 2016). Critically, Rosenkranz et al. (2021) demonstrated that neural synchrony during auditory narrative listening only occurs for the specifically attended narrative in competing-speaker scenarios. This suggests that selective attention is a mandatory filter during auditory listening. This built upon prior work by Dmochowski et al. (2012), who found that EEG-ISC was higher for intact versus temporally scrambled film excerpts, and that a narratively intact but non-engaging stimulus elicited lower ISC than critically acclaimed excerpts.

Audio stimuli have often been compared by utilizing “scrambling” methods to investigate the impact of intact stimuli. Stimuli are made incoherent by temporally reordering segments, which reduces the overall ISC. However, in audio-only contexts, when scrambled at the word level, ISC levels are similar between scrambled and unscrambled narrative stimuli (Ki et al., 2016); differences exist between spatial components, suggesting that some components may perhaps be sensitive to semantic processing rather than responsive to the auditory features of the stimulus and attentional state alone. Additionally, an intact, yet incomprehensible, stimulus (a narrative in Japanese presented to English speakers) was also included in Ki et al. (2016)’s work; participants showed reduced ISC to this incomprehensible narrative compared to the comprehensible narratives, which further supports the link between engagement and neural responses. Recent work has further demonstrated that EEG can track hierarchical linguistic features during continuous narrative comprehension (Broderick et al., 2018; Kaufeld et al., 2020), with neural oscillations organized into a content-specific hierarchy that reflects linguistic structure and meaning. To elucidate elements of semantic processing, it is therefore important to include a naturalistic semantic comparison, rather than a scrambled control or incomprehensible narrative. This points towards the utility of content-based, rather than temporally based, stimuli.

Despite these advances in understanding how the brain processes audio narratives, research has yet to systematically examine how specific semantic content dimensions, such as the social/non-social distinction, modulate EEG-ISC during auditory story listening. Auditory narratives have yet to be examined at the level of contextualized language, such as how certain language is used within a narrative, beyond the overall mood. While research has begun to identify which acoustic and semantic features of spoken stories, such as temporal intactness or comprehensibility of speech, drive EEG synchrony across listeners (Ki et al., 2016), less is known about how higher-level semantic content dimensions, such as social versus non-social information, modulate ISC. Given that fMRI research has demonstrated distinct neural processing for social content during naturalistic narrative processing (Yeshurun et al., 2017) and that social processing recruits specialized brain networks (Dodell-Feder et al., 2011; Saxe et al., 2004), examining whether social versus non-social content in audio narratives differentially affects EEG-ISC represents an important gap in the literature.

Thus, the current study aims to investigate the impact of overall content, examined through a combination of linguistic analysis, on neural engagement with auditory narratives. This results in the following research questions (RQs):

**RQ1** In naturalistic audio narratives, do social and non-social features of narratives impact overall EEG-ISC?
**RQ2** What is the relationship between self-reported behavioral engagement and EEG-ISC with naturalistic auditory narratives?

We aim to address these questions through content analysis of the narratives, EEG-ISC analyses, and participant responses to behavioral questions following each stimulus. For RQ1, we hypothesize that higher demands on social processing of text will elicit higher engagement based on evidence from fMRI research showing enhanced neural synchrony for social content (Yeshurun et al., 2017). For RQ2, we anticipate that higher levels of behavioral engagement will be correlated with higher levels of neural engagement, given our naturalistic comparison conditions and previous findings linking self-reported engagement to ISC during audio narrative listening (Dmochowski et al., 2012; Ki et al., 2016).

## 2 Methods

### 2.1 Ethics Approval Statement

The Stanford University Institutional Review Board (Protocol #50742) approved all procedures. All participants provided written informed consent prior to taking part in the study.

### 2.2 Participants

We recruited participants via the Stanford University participation pool. Requirements for participation included right-handedness, high fluency in English, normal hearing, aged between 18–35 years, and no history of recent concussion or diagnosed learning differences. Participants completed two experimental sessions, the first of which is the focus of this study. We continued data collection on a rolling basis until we reached 48 usable data sets; this was our target goal based on sample sizes of recent EEG-ISC studies involving stimuli of similar length (Kaneshiro et al., 2020; Dauer et al., 2021) and the unique counterbalancing design of the current experiment. We later excluded two participants during preprocessing due to significant noise artifacts across the data (*N* = 1) and an insufficient number of trials remaining after cleaning (*N* = 1), resulting in an analysis sample size of *N* = 46.

### 2.3 Stimuli

We selected six literary excerpt stimuli of varying genres. When possible, texts were chosen based on previously published literature (Kidd and Castano, 2013, 2019). The stimuli are summarized in Table 1; see Appendix A for opening and closing sentences of the excerpts.

**Table 1:** Literary excerpts and source. Title, author, excerpt source (as well as an approximation of genre), and duration for each stimulus.

| Title | Author | Excerpt Source (genre) | Duration (min:sec) |
|---|---|---|---|
| Uncle Rock | Dagoberto Gilb | Short story (contemporary fiction) | 4:06 |
| Nothing Living Lives Alone | Wendell Berry | Short story (historical fiction) | 4:33 |
| Corrie | Alice Munro | Short story (historical fiction) | 3:42 |
| Mountain Ways | Ursula K. Le Guin | Short story (science fiction) | 4:12 |
| How the Potato Changed the World | Charles C. Mann | Magazine article (nonfiction) | 4:23 |
| The Forest Unseen | David Haskell | Novel (literary science nonfiction) | 3:59 |

The selected excerpts were approximately 4 minutes long each (3:42–4:33) and consisted of approximately 600 words (571–618). The length of these stimuli was selected based on the length of previously presented stimuli in earlier ISC studies (e.g., Kaneshiro et al., 2020). We categorized each excerpt as either social or non-social by using two procedures. First, a qualitative team — comprised of the first and senior author — read through each text and placed the texts into two buckets based on the presence and role of character, world-building, dialogue, and plot. This led to the texts’ placement in a social or a non-social category. This process resulted in three texts that we categorized as social: the Gilb, Munro, and Berry texts. We classified the other three texts as non-social: the Le Guin, Mann, and Haskell texts. We refer to this distinction as the “condition”.

We later assessed the stimuli using LIWC (Boyd et al., 2022) and its built-in dictionary to evaluate vocabulary as a complementary means of categorization. The social texts contained more cognitive (e.g., “think”) and social words (e.g., “care”) on average, whereas the non-social texts used slightly more perception-related words (including spatial, e.g., “see” or “here”). One exception was the (non-social) Le Guin text which had a higher percentage of social words than the (social) Berry and about the same percentage of social words as the (social) Munro text. First, we explored all of these texts on a condition level to capture the overall quality as determined by the aforementioned qualitative team of human readers. We then computed percentages of social and cognitive words, which we used to establish a sociocognitive categorical spectrum (low, medium, and high), as analyzed in LIWC. The defining features of these texts are presented in Table 2 (by category) and Table 3 (by individual stimulus).

**Table 2:** Word counts and percentages per condition. Word count and LIWC dictionary word percentages per category.

| Condition | WC (sd) | Cognitive words in % (sd) | Social words in % (sd) | Perception words in % (sd) |
|---|---|---|---|---|
| S | 600 (20) | 11.29 (1.76) | 14.81 (4.75) | 10.17 (0.67) |
| NS | 589 (21) | 6.77 (1.22) | 7.95 (4.19) | 13.25 (0.84) |

**Table 3:**
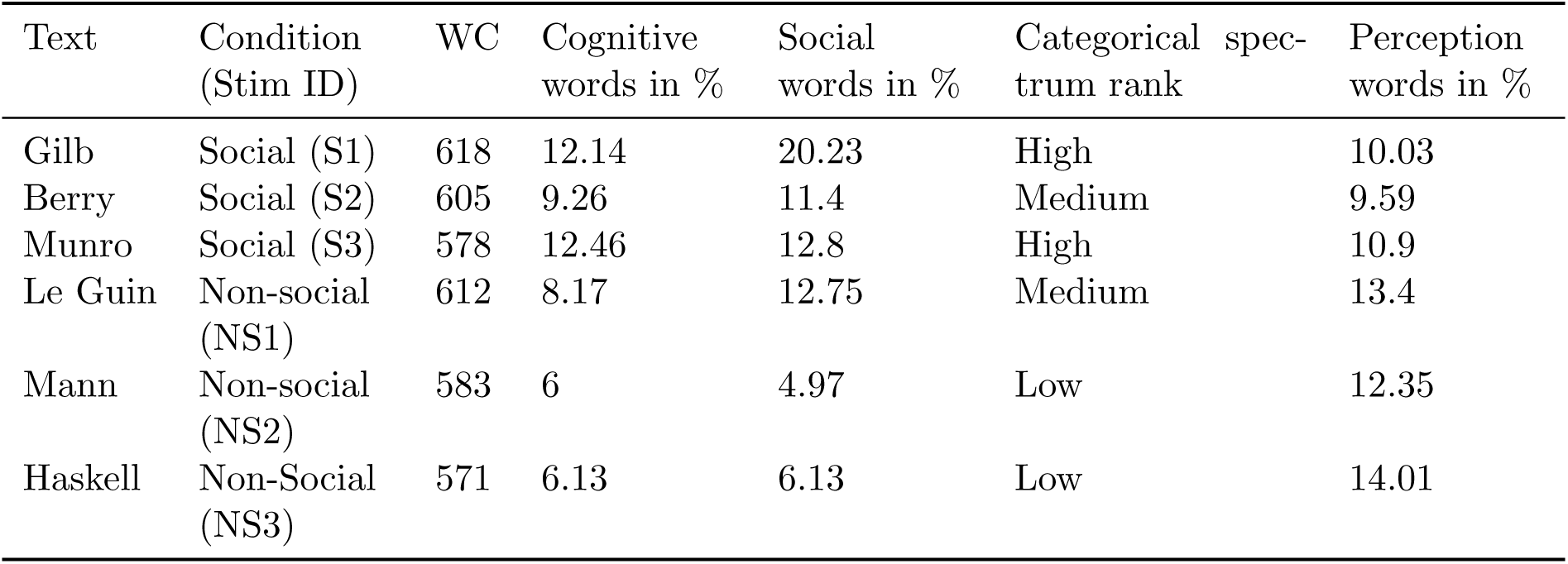
Stimulus characteristics and LIWC word percentages. Word count and LIWC dictionary word percentages per stimulus as well as categorical spectrum rank.

| Text | Condition (Stim ID) | WC | Cognitive words in % | Social words in % | Categorical spec-trum rank | Perception words in % |
|---|---|---|---|---|---|---|
| Gilb | Social (S1) | 618 | 12.14 | 20.23 | High | 10.03 |
| Berry | Social (S2) | 605 | 9.26 | 11.4 | Medium | 9.59 |
| Munro | Social (S3) | 578 | 12.46 | 12.8 | High | 10.9 |
| Le Guin | Non-social (NS1) | 612 | 8.17 | 12.75 | Medium | 13.4 |
| Mann | Non-social (NS2) | 583 | 6 | 4.97 | Low | 12.35 |
| Haskell | Non-Social (NS3) | 571 | 6.13 | 6.13 | Low | 14.01 |

Given our overarching goal to examine naturalistic stimuli, all excerpts were narrated by an adult female speaker. Recordings of these narrations were made using a voice-isolating Zoom H5 microphone; the files were trimmed in Audacity (https://github.com/audacity/ audacity) to remove silence at the beginning or end of the recordings and to remix the audio to mono. We further processed the .wav stimuli in MATLAB (https://www.mathworks.com) to present them during the experimental session by adding intermittent (every 1000ms) square-wave pulses; the function of these pulses was to establish regular timepoint stamping throughout stimulus presentation. The pulses were embedded in a separate audio channel and were not audible to participants.

### 2.4 Procedure

The experimental session lasted approximately 1.5 hours including set-up, breaks, and debriefing. We compensated all participants for their time. Participants completed demo-graphic information as well as questionnaires related to reading and music-listening habits during set-up and impedance checks in the experiment.

Participants were seated 1 meter away from a speaker located directly in front of them, which played the auditory stimuli. Each participant heard four of the six literary excerpt stimuli (two social and two non-social) per session to reduce participant burden, given the length of the session. The participant heard each literary excerpt only once, as previous research has shown that ISC is reduced when a stimulus is repeated (Dmochowski et al., 2012; Kaneshiro et al., 2020). Stimuli were assigned in a counterbalanced fashion across participants, ensuring equal assignment across the participant pool. Counterbalanced stimulus ordering further ensured that each stimulus was encountered equally in each possible presentation order, thereby mitigating potential effects of order and context.

The data analyzed here are part of a larger study comprising nine stimulus blocks: eight blocks of speech stimuli followed by one block of music stimuli. Each participant’s four assigned literary excerpt stimuli were distributed throughout the first eight blocks of the experiment such that one stimulus from each condition was presented in the first half of the experiment, and one of each condition in the second half. Furthermore, there was no chance for stimuli from the same condition to appear consecutively (e.g., a social excerpt could not appear in block 4 and block 5). Electrode impedance checks occurred before blocks 1, 5, and 9. An example participant session flow is presented in Table 4.

**Table 4:** Example Participant Array. Each participant completed nine experimental blocks total, including the four literary excerpt blocks analyzed here. Breaks followed each block, and impedance checks occurred before each of the three groupings reflected in the table rows. Blocks relevant to the current paper are in bold; other tasks are not the focus of the current manuscript. One excerpt was presented in each of the bolded blocks.

| Block | Condition |
|---|---|
| **1** | **Non-social literary stimulus** |
| **2** | **Social literary stimulus** |
| 3 | Other speech stimuli |
| 4 | Other speech stimuli |
| **5** | **Social literary stimulus** |
| 6 | Other speech stimuli |
| 7 | Other speech stimuli |
| **8** | **Non-social literary stimulus** |
| 9 | Music stimuli |

After the conclusion of each stimulus, participants answered three behavioral questions rating their interest and enjoyment, as well as how compelling they found the passage, on a 1–9 scale. These questions were always presented in the same order with the following wording:

1. How much of this passage was interesting, on a scale of 1 (none of it) to 9 (all of it)? Use the number keys at the top of the keyboard.
2. How compelling was this excerpt, on a scale of 1 (not compelling at all) to 9 (very compelling)? Use the number keys at the top of the keyboard.
3. How enjoyable was this excerpt, on a scale of 1 (not enjoyable at all) to 9 (very enjoyable)? Use the number keys at the top of the keyboard.

We adapted these subjective rating questions from previously published work (Kaneshiro et al., 2020; Dauer et al., 2021; Kaneshiro et al., 2024) so that engagement could be captured at both a neural and behavioral level.

### 2.5 EEG Acquisition and Preprocessing

We collected neural data using a 128-channel EEG net from the Electrical Geodesics, Inc. system (EGI; Tucker, 1993) with impedances below 50 kΩ. Impedances were checked both before and intermittently throughout data collection (see Table 4 for the timing of data impedance checks). We collected EEG data at a sampling rate of 1000 Hz, with reference to the vertex. We presented all components of the task using the Psychtoolbox toolkit in MATLAB (Kleiner et al., 2007), including audio stimuli and key-press response acquisition. Stimulus onset, as well as key-press response, was recorded using triggers sent to MATLAB. Continuous EEG recordings (one per block, including all vignette listening and participant responses) were exported to a MATLAB-compatible file format (.mat) using the inbuilt Net Station software. Preprocessing was then performed in MATLAB using custom, publicly available code (Malave et al., 2026); the procedure closely follows the cleaning process described in Kaneshiro et al. (2024) and is described in further detail there.

Data were preprocessed in combined, consecutive groupings of blocks that shared the same impedance check (blocks 1–4, blocks 5–8, and block 9). Each recording was high-pass filtered (0.3 Hz), notch filtered (59–61 Hz), and low-pass filtered (50 Hz). EEG data were downsampled by a factor of 4 to an analysis sampling rate of 250 Hz. The data were then epoched to the length of each stimulus. Following epoching, preprocessing continued by examining potential bad channels and identifying any channels that were significantly noisy or uncorrelated from neighboring electrodes. These channels were then temporarily removed from the data frame. We also computed electroculogram (EOG) channels from electrodes surrounding the eyes (above and/or below for VEOG, on the side of HEOG).

We continued to preprocess the data using extended Infomax Independent Components Analysis (ICA) to remove artifacts associated with eye movement, ECG, and DIN (digital input trigger) artifacts. This removal was done in a semi-automatic process using the EEGLAB ICA function (Delorme and Makeig, 2004), as incorporated into the broader pipeline (Malave et al., 2026).

After ICA, we temporarily removed from each trial any remaining electrodes with 10% or more of voltage magnitudes greater than 50 *µ*V. Median-based direct current (DC) correction was performed on data within a trial, and data points exceeding four standard deviations of the mean power were replaced with NaNs; this process was performed on a per-electrode basis and iterated through in four stages. Matrix rows representing bad electrodes were then reintroduced into the data as NaNs, along with a row of zeros that represented the Cz reference. Then, the data were converted to average reference, ignoring NaNs. Lastly, we imputed all NaNs, including those representing previously removed bad channels, by computing spatial averages from the data of its neighbor sensors, followed by a mean-based DC correction.

During this process, we removed two participants: one participant was removed due to gross noise artifacts for two trials across the literary stimuli (resulting in only 50% of the relevant data remaining), and one participant had significant noise artifacts across the entirety of their data set.

After preprocessing, we aggregated the trials into one 3D electrode-by-time-by-participant data matrix for each stimulus. As each participant experienced only 4 of 6 available stimuli, the final participant counts for each stimuli are lower than the overall number of participants. The final size of the matrices are in Table 5 below.

**Table 5:** Stimuli Data Matrices. 3D data matrices per stimulus.

| Text | Stimulus ID | Final Matrix Dimensions (electrode x time x participant) |
|---|---|---|
| Gilb | S1 | 125 × 61550 × 30 |
| Berry | S2 | 125 × 68326 × 30 |
| Munro | S3 | 125 × 55612 × 29 |
| Le Guin | NS1 | 125 × 62987 × 30 |
| Mann | NS2 | 125 × 65699 × 32 |
| Haskell | NS3 | 125 × 59822 × 30 |

### 2.6 Data Analysis

For analyses, we used cleaned EEG responses to social or non-social vignettes, alongside their behavioral responses. We first spatially filtered the EEG data using Reliable Components Analysis (RCA), and then computed the ISC separately for each stimulus. ISC values represent a summative measure of the correlation of EEG data across subjects over the entirety of a given stimulus, and is calculated separately for each Reliable Component (RC) derived from the earlier RCA step. We then used the distributions of ISC values to compare and analyze the responses for engagement per stimulus and per overall condition.

#### 2.6.1 EEG Spatial Filtering

We used the approach developed by Dmochowski et al. (2012) to spatially filter our data prior to calculating the ISC. This step, which uses an generalized eigenvalue decomposition process called Reliable Components Analysis (RCA), increases the signal-to-noise ratio (SNR) of the data while reducing the spatial dimensionality of the data from over 100 sensors to a small number of spatial components (Dmochowski et al., 2012). As this approach maximizes the across-trials covariance (to a stimulus) relative to the within-trials covariance, it produces spatial components with maximally correlated activity across trials. The spatial components, or RCs, are calculated in order of how much reliability within the data is explained (with RC1 being the most, RC2 being the second-most, and so forth), and represent linear combinations of the electrodes.

RCA was calculated separately for social and non-social stimuli (three stimuli per RCA run) using publicly shared code for MATLAB (Dmochowski et al., 2015). Although ISC research varies in the number of components considered, we began by computing the top five RCs. In our categorical level calculations, RC1, RC2, and RC3 were significant across both the social and non-social stimuli (all p *<* 0.001, permutation test). RC4 and RC5 were not consistently significant across conditions, and thus were not used for subsequent analyses. This is in line with previous research, which tends to focus on either RC1 alone or the first three components (e.g., Ki et al., 2016; Rosenkranz et al., 2021; Ueno and Shimada, 2023). Visualized topographies of the RCs represent forward-model projections of the component weights (Parra et al., 2005).

#### 2.6.2 Inter-Subject Correlation

ISC calculations were performed on a per-stimulus, per-RC basis using a one-against-all approach such that, across the duration of each stimulus, each response vector was correlated with every other response vector within a given component. These values could then be averaged across participants to obtain one ISC value per participant for each stimulus and component. Following previous work (e.g., Ki et al., 2016), ISC could further be summed across the three RCs to produce a single per-participant value for a given stimulus.

### 2.7 Statistical Testing

Our statistical analyses were completed with a combination of the MATLAB Psychophysics Toolbox (Brainard, 1997) and R (RStudio; Posit Team, 2025). In MATLAB, the significance of the RCA results per-component were calculated using permutation testing, using previously established methods (Dauer et al., 2021; Kaneshiro et al., 2020). Here, each stimulus underwent 1,000 iterations in which the phase spectrum of each EEG trial was independently randomized prior to RCA. We then used the resulting distribution of 1,000 outcomes as a null distribution. Comparing a null distribution to the observed data allowed us to compute p-values for each RC for each stimulus, as well as each condition.

Following our permutation and significance testing, the remaining statistical analyses were completed using R. A complete list of used packages can be provided upon request. We started by examining how ISC changes when there are multiple stimuli within one category; thus, we reviewed our data for the impact of presentation order (whether it was the first, second, third, or fourth stimulus) on ISC. This is an important manipulation given past research in ISC has suggested that repeating a single stimulus reduces ISC and neural responses more broadly (e.g., Grill-Spector et al., 2006; Nastase et al., 2019).

For our behavioral ratings, we began by checking the distributions of each question as well as the sum of all three questions added together. Following previous research (Dauer et al., 2021; Kaneshiro et al., 2020), our ordinal Likert scale responses were handled as approximately continuous for statistical purposes (Norman, 2010). We thus calculated the mean across all responses, as well as within each condition, for individual questions and their sum. We used the Wilcoxon signed-rank test given the non-normal distribution of the data and to analyze whether or not there were significant differences in behavioral responses based on both condition or individual stimuli.

We then used the lme4 package (Bates et al., 2015) to examine the impact of condition, as well as behavioral ratings, on ISC. We describe each model below; models 1 and 1.5 address RQ1. Model 2 addresses RQ2, and model 3 examines any potential interaction between these two RQs.

We performed a linear mixed-effects model analysis of the impact of condition on ISC first by running our model with the fixed effect of condition, and the random effects of stimulus and participant (Model 1). We repeated this model for each component (RC1, RC2, and RC3) as well as the sum of all 3 components. Each model was then compared against a null model, without the fixed effect of condition, using the lme4 anova function, resulting in a chi-squared value and p-value for each model. We then repeated this model and process (Model 1.5) using spectrum ranking (high, medium, or low) as a related metric of condition. To examine the impact of behavioral responses on ISC, we repeated the above process with fixed effects of behavioral response (summed across all 3 questions), and random effects of stimulus and participant (Model 2). Each model was then compared against a null model without the fixed effect of behavior. Lastly, we checked for an interaction between condition, as well as spectrum ranking, and behavioral ratings by adding condition as an interactor (Model 3 and Model 3.5). Each model was compared against a null model with fixed effects of condition and behavior and no interaction effect.

For these analyses, which involved multiple comparisons, we corrected our results using the False Discovery Rate (FDR; Benjamini et al., 2001) adjustment on the necessary p-values, as has been done in previous ISC work using linear mixed effects models (e.g., Dauer et al., 2021; Kaneshiro et al., 2020).

## 3 Results

To explore our primary research questions, we begin by reviewing our behavioral findings and ISC results at a descriptive level. We then examine the impact of condition and spectrum ranking on ISC for all six stimuli, repeating these same steps for exploratory analyses on a subset of the data.

### 3.1 Behavioral

After listening to each stimulus, participants responded on a Likert scale of 1–9 about how much they enjoyed, were compelled by, and were interested in what they had just heard. We also summed those values. These responses are represented at a stimulus level in Table 6 below. At the conditional level, the mean of the social category was 5.18 (sd = 2.06) and the mean of the non-social category was 5.05 (sd = 2.06).

**Table 6:**
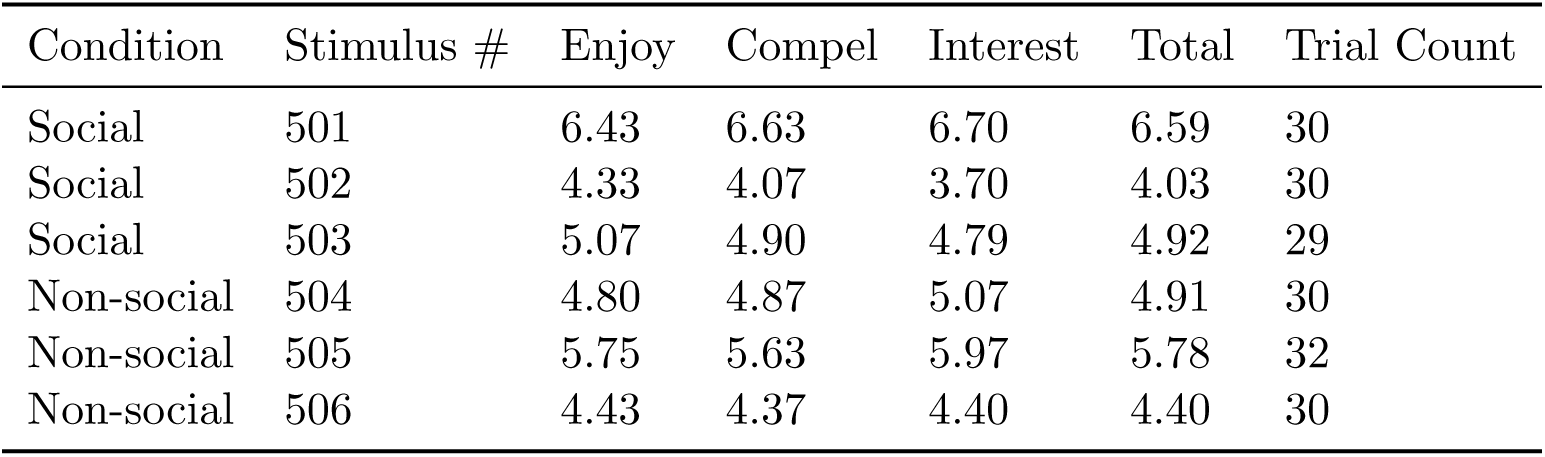
Behavioral Ratings per Stimulus. Behavioral ratings by stimuli, with the mean of sum of all three shown as the total. Trial counts are also displayed given that each participant only saw 4 out of the 6 available stimuli.

| Condition | Stimulus # | Enjoy | Compel | Interest | Total | Trial Count |
|---|---|---|---|---|---|---|
| Social | 501 | 6.43 | 6.63 | 6.70 | 6.59 | 30 |
| Social | 502 | 4.33 | 4.07 | 3.70 | 4.03 | 30 |
| Social | 503 | 5.07 | 4.90 | 4.79 | 4.92 | 29 |
| Non-social | 504 | 4.80 | 4.87 | 5.07 | 4.91 | 30 |
| Non-social | 505 | 5.75 | 5.63 | 5.97 | 5.78 | 32 |
| Non-social | 506 | 4.43 | 4.37 | 4.40 | 4.40 | 30 |

The distributions for each of these responses, as well as their total, were found to be non-normal (as expected of Likert data; Norman, 2010). Given the non-normal distribution, we examined whether or not condition (social or non-social) was significantly related to each question. We found no significant differences between conditions on any of the three questions or their total (all p *>* 0.4). At the stimulus level, we found some significant differences between certain stimuli for each of the questions, as well as their total. These are summarized below in Table 7, with the adjusted p-value for multiple comparisons (utilizing the Holm-Bonferroni method; Holm, 1979). Table 6 contains the means of the distributions being compared in Table 7.

**Table 7:**
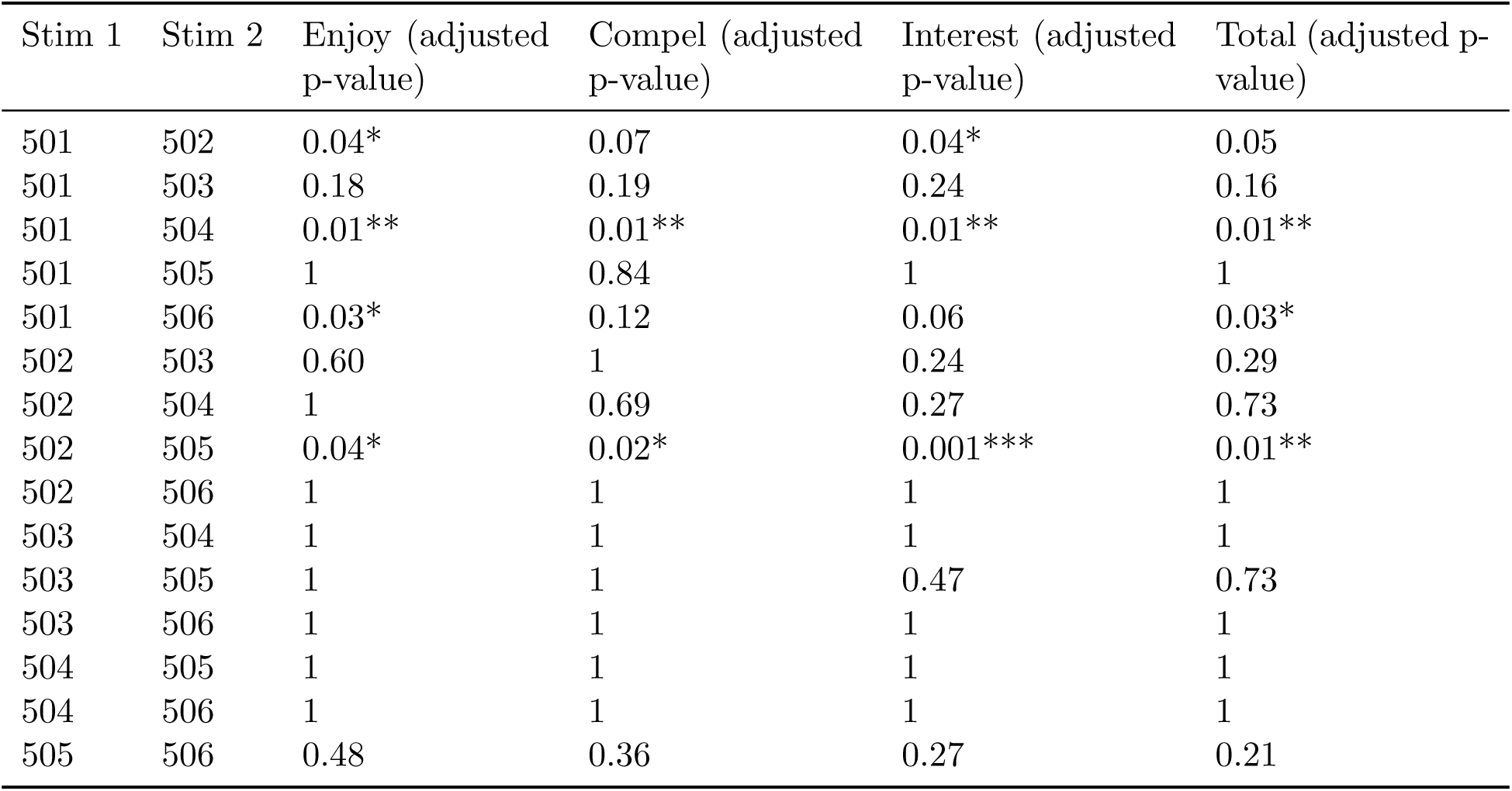
Behavioral Question Pairwise Comparisons. Pairwise (two stimuli per pair) comparisons using the Wilcoxon signed-rank test to examine significant differences per behavioral question (and their total). Adjusted p-values were computed using the Holm-Bonferroni method. Significance levels: *p*<*0.05, **p*<*0.01, ***p*<*0.001.

| Stim 1 | Stim 2 | Enjoy (adjusted p-value) | Compel (adjusted p-value) | Interest (adjusted p-value) | Total (adjusted p-value) |
|---|---|---|---|---|---|
| 501 | 502 | 0.04* | 0.07 | 0.04* | 0.05 |
| 501 | 503 | 0.18 | 0.19 | 0.24 | 0.16 |
| 501 | 504 | 0.01** | 0.01** | 0.01** | 0.01** |
| 501 | 505 | 1 | 0.84 | 1 | 1 |
| 501 | 506 | 0.03* | 0.12 | 0.06 | 0.03* |
| 502 | 503 | 0.60 | 1 | 0.24 | 0.29 |
| 502 | 504 | 1 | 0.69 | 0.27 | 0.73 |
| 502 | 505 | 0.04* | 0.02* | 0.001*** | 0.01** |
| 502 | 506 | 1 | 1 | 1 | 1 |
| 503 | 504 | 1 | 1 | 1 | 1 |
| 503 | 505 | 1 | 1 | 0.47 | 0.73 |
| 503 | 506 | 1 | 1 | 1 | 1 |
| 504 | 505 | 1 | 1 | 1 | 1 |
| 504 | 506 | 1 | 1 | 1 | 1 |
| 505 | 506 | 0.48 | 0.36 | 0.27 | 0.21 |

The most significant contrasts were in the pairs of stimuli 501-504, 502-505, and 501-506. These were largely driven by higher behavioral scores for 501 (social) and 505 (non-social).

### 3.2 Reliable Spatial Components

For our ISC analyses, we began by spatially filtering the data for each stimulus condition to reduce the data from 125 electrodes to its maximally correlated reliable components (producing eigenvectors which serve as the spatial filters). RCA also produces a vector of corresponding eigenvalues (referred to here as dGen values) that convey the overall correlation of each component. In Figure 1, we present the social and non-social dGen values with their value and significance level, as assessed via permutation testing.

**Figure 1:**
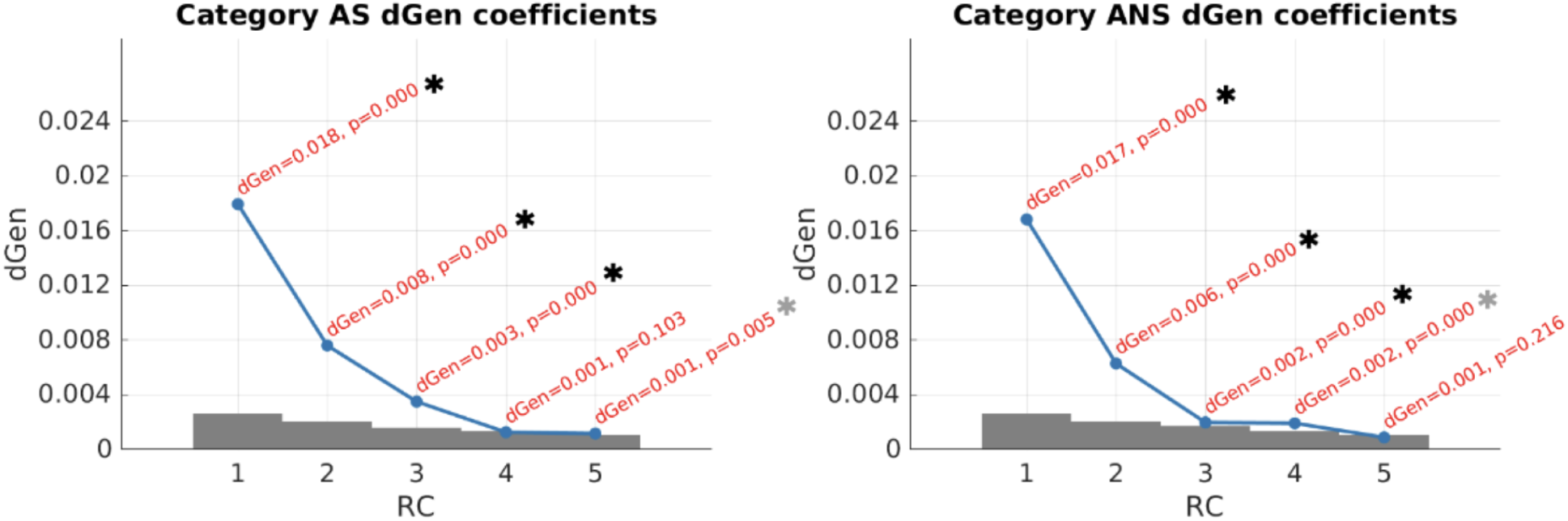
Categorical dGen Coefficients. The eigenvalues (generated during RCA) that serve as correlation coefficients, along with their associated significance levels. Reliable components (RCs) that are significant for both conditions are marked with a black asterisk; RCs that are only significant for one condition are marked with a gray asterisk. The gray bars underlying the dGen line represent the null distribution’s 95th percentile as calculated during the permutations during RCA.

We find that RC1, RC2, and RC3 are statistically significant for both conditions. Although RC4 is significant in the non-social category, it is not in the social category; similarly, RC5 is significant for the social category and not the non-social category. Because of this, we focus only on components 1, 2, and 3 for the remainder of our analyses. In Figure 2, we show the spatial components (via topography) for RC1, RC2, and RC3. We do not report RC4 and RC5 as these RCs and their associated correlation coefficients were not reliably significant across categories and stimuli.

**Figure 2:**
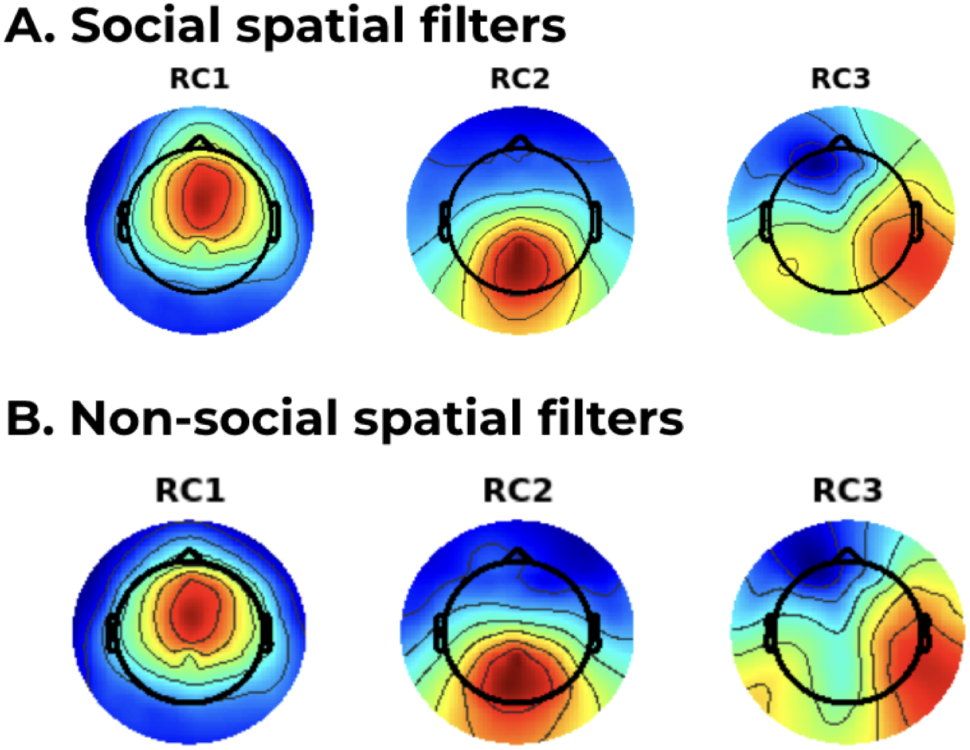
Topographies (Spatial Filters) Per Category. The spatial filter weights by category for social (A) and non-social (B) across RC1, RC2, and RC3. Polarities of the topographies were reversed as needed for simplified visual interpretation; this does not impact ISC calculations.

Additionally, we computed Pearson correlations between conditions per component (e.g., RC1 of social and RC1 of non-social). All three components were strongly and statistically significantly correlated with their counterparts in the other condition (r *>* 0.96, p *<* 0.001).

### 3.3 Inter-Subject Correlation

#### 3.3.1 RQ1: Sociocognitive Group Impacts ISC in RC3

As earlier noted, we focused our statistical modeling on RC1, RC2, RC3, and the sum of RC1 through 3 (aggregate RC) due to inconsistent significance in RC4 and RC5 (Figure 1). We hypothesized that higher demands of sociocognitive processing would result in greater ISC due to increased engagement. We hypothesized that this might be most evident in RC2 or RC3, given that previous research has shown that RC1 is more reflective of attentional engagement rather than content.

We examined the impact of content through two variables: condition (a qualitative judgment by authors of this study) and sociocognitive ranking (a quantization of the number of sociocognitive words into low, medium, and high). We modeled the fixed effect of condition or sociocognitive ranking on ISC, with random effects of participant and stimulus; our null model excluded the fixed effect. We corrected for multiple comparisons using the FDR method (*α* = 0.05). After adjustment, we found that there was no significant relationship between condition and ISC for RC1, RC2, RC3, or the RC sum. For sociocognitive ranking, pairwise comparisons revealed no significant difference between the low and medium groups (b = 6.21e-05, SE = 0.00102, t(2.97) = -0.06, p = 0.956). However, significant differences were found between low and high groups (b = -4.36e-03, SE = 0.00102, t(2.94) = 4.28, p = 0.037) and between medium and high groups (b = -4.42e-03, SE = 0.00103, t(3.05) = -4.30, p = 0.037). Although the contrast between low and medium was not significant, we still found that it followed approximately the same positive trend from low to high cognitive group, which we represent graphically in Figure 3.

**Figure 3:**
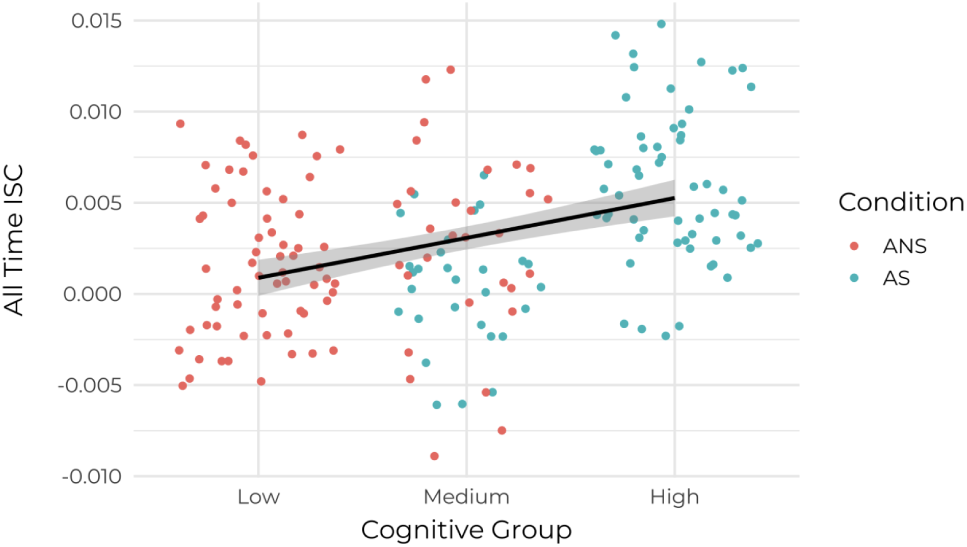
Correlation of ISC and Cognitive Group. A scatterplot of inter-subject correlation (ISC) scores for RC3 by sociocognitive group (low, medium, and high). Individual data points are shown per participant, with the social category in blue and the non-social category in red. The linear regression line (black) and 95% confidence interval standard error (gray) represent the general positive trend between increasing sociocognitive word usage and ISC.

#### 3.3.2 RQ2: Behavioral Ratings Are Reflected in RC2 and Aggregate RC ISC

We hypothesized that higher levels of behavioral engagement would be correlated with higher ISC values. We hypothesized that this effect might be most prevalent in RC1 or RC2, given conflicting evidence about which of these components may be related to ISC in the literature.

Given earlier findings that individual questions did not differ significantly across condition or sociocognitive group ranking, we proceeded with the mean score of all three behavioral questions together (interest, compulsion, and enjoyment), which we will call the “Aggregate Behavioral Response”. We modeled this as a fixed effect on ISC, with random effects of subject and stimulus. After FDR correction for multiple comparisons (*α* = 0.05), significant positive associations were found for RC2 (*β* = 5.77e-04, SE = 1.69e-04, t(168) = 3.41, p = 0.004) and aggregate RC (*β* = 1.09e-03, SE = 3.48e-04, t(166) = 3.12, p = 0.004). No significant relationships were observed for RC1 (*β* = 2.16e-04, SE = 2.10e-04, t(168) = 1.02, p = 0.428) or RC3 (*β* = 3.53e-06, SE = 1.58e-04, t(134) = 0.02, p = 0.957). We represent these findings in Figure 4, with RC2 in panel A and the aggregate RC in panel B.

**Figure 4:**
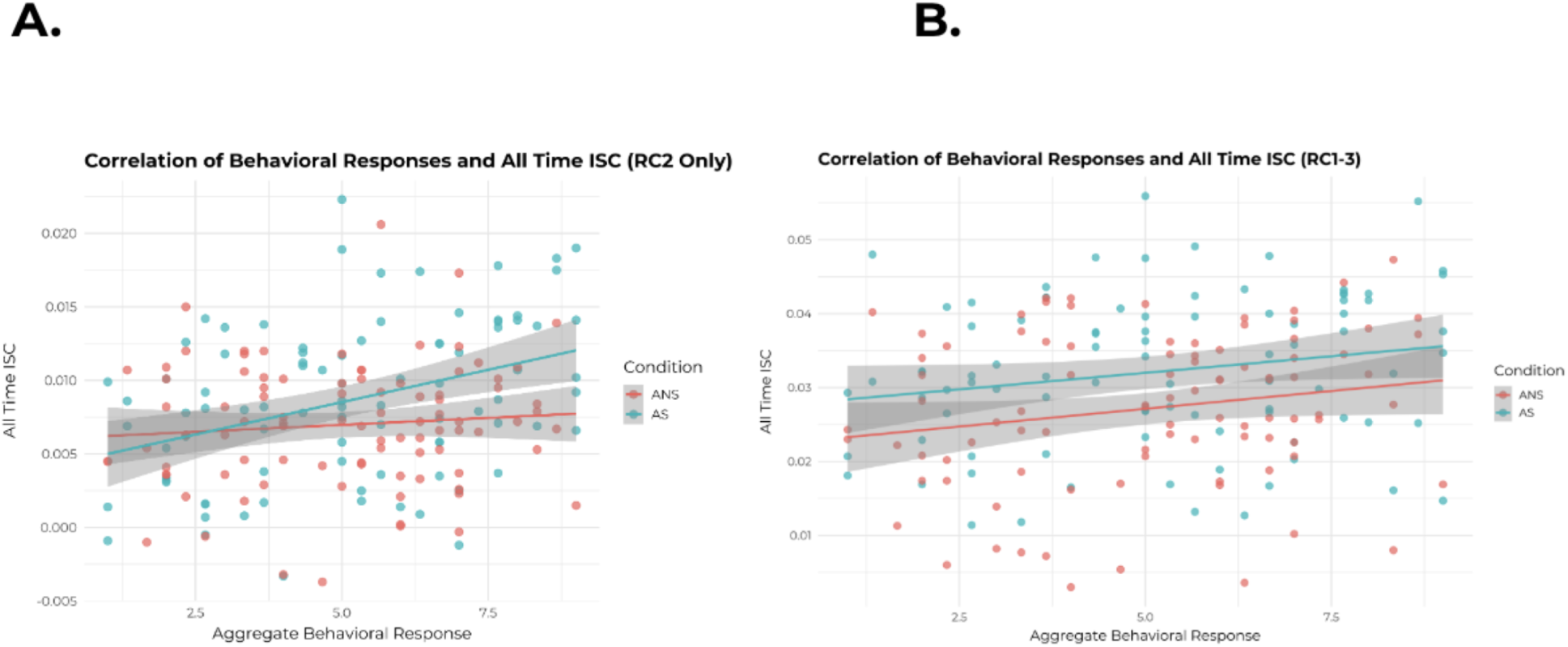
Correlation of ISC with RC2 and Aggregate RC. A scatterplot of inter-subject correlation (ISC) scores for RC2 (panel A) and aggregate RC (panel B) by aggregate behavioral response. Individual data points are shown per participant, with the social category in blue and the non-social category in red. The linear regression line (in correlating colors per category) and 95% confidence interval standard error (gray) represent the general positive trend between increasing behavioral response and ISC in RC2 and aggregate RC.

#### 3.3.3 Disentangling Interaction Versus Main Effects

Given the representation of the data in Figure 4 (panel A), we also examined our data for an interaction effect between condition (as well as sociocognitive ranking) and behavioral response. Using an ANOVA, we compared this model (which included random effects of subject and stimulus) against a null model which included behavioral response and condition as fixed effects only.

After FDR correction on our ANOVA comparisons, we did not find a significant interaction between behavioral response and condition on ISC (all FDR-adjusted p *>* 0.10). Below, we discuss some of our modeling of behavior and response condition as separate fixed effects (rather than an interaction). We interpret these results cautiously given lower degrees of freedom for some of these comparisons.

However, in our aggregate RC null model, we saw that there was a significant impact of both behavioral response and condition (as fixed effects) on ISC. Both behavioral scores, *β* = 1.21 × 10*^−^*^3^, SE = 3.21 × 10*^−^*^4^, t(60) = 3.77, p *<* 0.001, and condition, *β* = 4.72 × 10*^−^*^3^, SE = 1.25 × 10*^−^*^3^, t(2.89) = 3.77, p = 0.035, significantly predicted ISC.

We also repeated this modeling process with our sociocognitive ranking variable. Again, we found no significant interactions between behavioral response and sociocognitive ranking on ISC (all FDR-adjusted p *>* 0.10). However, we once again found an additive impact of our two fixed effects in the null models, this time in RC2 and aggregate RC. For RC2, aggregate behavioral response significantly predicted ISC (*β* = 5.59e-04, SE = 1.65e-04, t(107) = 3.40, p *<* 0.001), with higher scores associated with larger ISC values. Additionally, the high sociocognitive group showed slightly higher ISC compared to the low group (*β* = 2.53e-03, SE = 7.75e-04, t(2.56) = 3.27, p = 0.058). Similarly, for aggregate RC, behavior positively predicted ISC (*β* = 1.08e-03, SE = 3.48e-04, t(151) = 3.11, p = 0.002), and the high sociocognitive group exhibited marginally larger ISC values than the low group (*β* = 5.53e-03, SE = 2.17e-03, t(2.78) = 2.55, p = 0.091).

These results suggest that in RC2 and aggregate RC, ISC scores are independently impacted by both the content of the passage (condition for aggregate RC, sociocognitive ranking for RC2 and aggregate RC) as well as the behavioral responses of the listener. Thus, linear mixed effects modeling that accounts for both content and behavioral engagement may better predict the related ISC values, as higher behavioral responses as well as greater social/cognitive content trend towards higher ISC.

#### 3.3.4 Impact of Stimulus Order

Lastly, we analyzed our ISC values for the impact of order effects as a methodological contribution for better understanding ISC. This analysis will not impact our statistical testing, given the counterbalanced nature of our literary stimuli across the entire experiment. However, although we counterbalanced and did not repeat individual stimuli with our participants, we repeated conditionally similar stimuli (2 social, 2 non-social) per single experiment; all four stimuli also had highly similar narrative structures. Given previous research that repeated stimuli may reduce ISC values, we examined the impact of order within condition (first or second) and overall presentation (first, second, third, or fourth). We did so by running a linear mixed effects model with the fixed effects of either condition presentation or overall presentation on RC1 and the sum of RC1 through 3 (with random effects for stimulus and participant). After applying FDR for multiple comparison corrections, we found no significant differences for condition order (p *>* 0.1). However, we did find that overall presentation order marginally impacted ISC (p = 0.08) such that there was a slight decrease in ISC value by the end of the experiment.

## 4 Discussion

In summary, this study of naturalistic audio narrative disentangles three dissociable contributors to RC-related effects in ISC while also emphasizing the consistency and utility of ISC as a neural metric of engagement with a stimulus.

This study’s topographic findings highlight that reliable components are stable across content-related conditions in narrative processing. Given current knowledge on RCs in EEG-ISC research, our results further support that RC1 likely captures a combination of auditory and attentional processing that is stimulus-driven and consistent across participant groups (Dmochowski et al., 2012). This interpretation aligns with the hierarchical timescale framework established in narrative comprehension research, wherein auditory cortical regions synchronize rapidly to acoustic features while higher-order regions engage over longer timescales to process meaning (Lerner et al., 2011; Wilson et al., 2008). We novelly extend previous research in ISC by examining the role of engagement in RC2 and the role of sociocognitive content in RC3 in relation to naturalistically content-manipulated audio narratives (rather than narratives manipulated by scrambling or language). Our findings suggest that RC2 appears to be sensitive to behavioral engagement, whereas RC3 is sensitive to content.

Our condition-related finding suggests that higher social-related processing increases ISC in RC3. This increase aligns with previous research in reading that suggests that, when in possession of similar neural resources, readers interpret sociocognitively-involved passages more similarly to one another (e.g., Van den Broek and Helder, 2017), which results in higher ISC values. This finding extends the framework proposed by Nguyen et al. (2019), who demonstrated that shared understanding of narrative meaning drives neural synchrony. Our results suggest that this principle operates specifically for socially demanding content, and manifests in distinct neural components. The content-sensitivity of RC3 is also consistent with work showing that EEG tracks hierarchical linguistic features during narrative comprehension, with content-specific hierarchies that reflect meaning-level processing (Broderick et al., 2018; Kaufeld et al., 2020).

Our behavioral-related finding validates previous research that suggests that ISC reflects meaningful processing at a neural level, with higher levels of engagement and attention connected to higher ISC values (e.g., Kaneshiro et al., 2024). This relationship between behavioral engagement and RC2 is consistent with Cohen (2018)’s proposal that high ISC is a biological marker of narrative engagement, wherein engaged listeners’ brain activity drives a shared perception of a stimulus. The specificity of this effect to RC2 (and subsequently aggregate RC), suggests that this engagement may be specifically related to the level of processing that is captured by RC2. This extends previous work in audio-only narrative ISC research that demonstrated links between behavioral engagement and neural synchrony (Ki et al., 2016) by identifying RC2 as the specific component that may underlie this relationship. This dissociation between behavioral engagement effects (RC2) and content effects (RC3) provides empirical support for theoretical models proposing that ISC reflects multiple states of neural alignment, from basic attentional filtering (as demonstrated in a competing-speaker paradigm; Rosenkranz et al., 2021) to higher-level semantic integration (Cohen and Parra, 2016). The lack of interaction between these two findings extend previous research that ISC is both stimulus and state driven. Instead, both the content of the stimulus as well as the behavioral engagement of the listener determine neural engagement with naturalistic stimuli. This dual influence mirrors the coupled neural systems framework established in spoken narrative research, wherein successful comprehension requires both stimulus-driven synchrony and engagement-related processes (bottom-up and top-down; Silbert et al., 2014; Stephens et al., 2010). Intuitively, a listener may be engaged with the task behaviorally but may be less interested by vocabulary related to fictional character interaction, as they instead prefer texts focused on perceptual and scientific knowledge. This, alongside other state traits such as tiredness or place in the experiment, are important individual differences that may factor into ISC as a measure. Individual variability in neural responses to narratives is well-documented, with ISC patterns showing sensitivity to age, sex, and internal representation differences (Petroni et al., 2018), suggesting that even within engaged listeners, personal relevance and comprehension strategies may modulate neural synchrony.

Taken together, our results also have multiple implications for ISC research in EEG. Our component-specific findings suggest that neural components can be dissociated by function; this provides support for earlier theoretical proposals that ISC reflects multiple states of neural alignment, from processes generally considered low-level (such as auditory processing) to processes considered high-level (such as semantic meaning-making; Cohen and Parra, 2016). Specifically, our findings that distinct RCs track different narrative features align with the hierarchical processing architecture demonstrated in fMRI work, wherein slow cortical dynamics at different timescales accumulate information from immediate acoustics and extended narrative arcs (Honey et al., 2012; Simony et al., 2016). The EEG-based components may represent electrophysiological signatures of these hierarchical processes, offering temporal resolution complementary to the spatial precision of fMRI-based narrative studies using audio narratives. The component-level differentiation we observe in EEG-ISC may reflect the electrophysiological counterpart to the network segregation observed in fMRI work (e.g., Saxe et al., 2004; Yeshurun et al., 2017).

Additionally, we find that ISC captures both objective (sociocognitive word count) and subjective (behavioral ratings) measures of engagement, supporting previous work that suggests that ISC during narrative comprehension reflects multiple levels of processing that encompass both stimulus features and engagement (Simony et al., 2016). Furthermore, the sensitivity of different components to semantic content versus self-reported engagement parallels findings that shared neural responses can differentiate between conscious processing states (as demonstrated in clinical populations with consciousness disorders; Iotzov et al., 2017), suggesting that component-level ISC analysis may reveal dissociable social processes operating during comprehension.

Lastly, our finding regarding order effects (and the role of overall presentation, not within-condition order) suggests that ISC may be subject to global fatigue effects (Nastase et al., 2019), rather than simple stimulus repetition, and has important implications for future ISC experimental designs. These order effects may reflect changes in the brain’s capacity to maintain the long-timescale information accumulation required for narrative comprehension (Honey et al., 2012), with fatigue potentially disrupting the sustained neural activity necessary for deep engagement with narratives.

### 4.1 Limitations and Future Work

It should be noted that, in the current study, our participant group is highly homogeneous. Previous research has suggested that shared participant features drive ISC (such as personality traits or demographics, e.g., Finn et al., 2018; Lahnakoski et al., 2014), and so these findings should also be replicated with more diverse participant groups. There were also no clear measures of individual differences that might have addressed questions about preference — whether someone prefers a non-fiction piece over another type of text, for example — that may have helped us understand participants who did not fit the general trend of higher ISC for social texts. Future work might incorporate measures of reading preferences, prior knowledge, or theory of mind abilities, as individual differences in social cognition could modulate neural synchrony in socially rich narratives similarly as to how comprehension similarity drives ISC (Nguyen et al., 2019). Additionally, the current study uses a retrospective behavioral measure, rather than a moment-to-moment behavioral probes. Other recent EEG-ISC studies (e.g., Dauer et al., 2021; Kaneshiro et al., 2024) have used measures of continuous behavior that may address this; we may also seek to explore moment-to-moment engagement through additional analytic approaches, such as time-resolved ISC (Dmochowski et al., 2012; Poulsen et al., 2017; Dauer et al., 2021; Kaneshiro et al., 2024).

Time-resolved work may answer this question by looking at moment-to-moment ISC, as aligned with the timing of the narrative, and examining neural engagement at critical narrative moments. Such analyses would align with the timescale-specific processing network (Lerner et al., 2011) and could reveal how different RCs track narrative features that unfold over different temporal windows, from word-level semantic processing captured by RC3 to sustained attention reflected in RC2. This may highlight whether specific narrative moments that involve social content, such as character interactions, drive neural synchrony more than other moments. Time-resolved ISC could also test whether anticipatory neural coupling (similar to Stephens et al., 2010) predicts comprehension of upcoming events. Additional further work may also look at behavioral responses as predictors (e.g., Cohen et al., 2017) to improve our understanding of the potential direction of the relationship between ISC, naturalistic narrative, and behavioral response. Moreover, examining ISC’s predictive validity for downstream outcomes (such as memory of narrative content; e.g., Cohen and Parra, 2016) would establish whether component-specific synchrony during auditory story listening predicts long-term retention. This approach would be particularly relevant for educational applications given evidence that student-teacher brain synchrony predicts learning outcomes in lecture contexts (Davidesco et al., 2023).

## 5 Conclusion

Overall, the findings of the current study suggest that different reliable components (RCs) can be used to capture different levels of processing within a broader hierarchy (Lerner et al., 2011). RC1 was not significantly related to either content or behavior, enforcing that it may be related to low-level attention and sensory processing. This pattern is consistent with the resilience of basic auditory synchrony even under challenging listening conditions (e.g., Irsik et al., 2022), and also reflects the mandatory attentional filtering required for selective narrative engagement (Ki et al., 2016). RC2 was significantly related to behavioral responses, establishing a brain-behavior connection across the dimensions of interest, compulsion, and enjoyment. This finding positions RC2 as potentially reflecting the engagement-related neural processes that enable shared perception during emotionally salient narratives (Cohen, 2018), consistent with supramodal processing that operates above basic sensory features and extends previous audio ISC research involving states of engagement and ISC (Ki et al., 2016).

Lastly, RC3 was related to the content of the narrative, suggesting a relationship to semantic and sociocognitive processing. The content-sensitivity of RC3 aligns with the electrophysiological tracking of semantic dissimilarity and linguistic structure during natural speech comprehension (Broderick et al., 2018; Kaufeld et al., 2020) and may reflect the integrative neural processes that are required to construct coherent narrative representations across timescales (Honey et al., 2012; Davidesco et al., 2023). These component-specific effects are novel in that they suggest utility in analyzing RC1, RC2, and RC3 separately, rather than focusing on RC1 and an aggregate RC score alone. ISC is not just a measurement of attentional engagement, but may also capture multiple simultaneous processes that are all necessary during naturalistic narrative processing. This multi-component framework for understanding ISC during auditory narrative comprehension offers a bridge between EEG-based investigations of engagement and fMRI studies of hierarchical narrative processing, while also providing an ecologically valid approach to studying how the brain constructs meaning from the continuous speech streams encountered in everyday contexts.

Furthermore, this study broke new ground by demonstrating what can be learned from the innovation of contrasting authentic literary narratives. By moving beyond scrambled or incomprehensive controls, and instead using excerpts from established literary works, we probed how the brain responded to nuanced, ecologically valid variations in content. This approach revealed that the synchronized neural response to stories is not monolithic, and that different components tracked dissociable aspects of the narrative experience. This multi-faceted view of neural engagement, facilitated by the component findings we have shared in this work, were made possible by our naturalistic stimulus design. They underscore the brain’s sophisticated capacity to simultaneously process form, content, and personal relevance of a story; overall, this work helps to advance our understanding of how we connect with the narratives that shape our worlds.

## Acknowledgments

The authors would like to thank Suanna Moron, Vani Dewan, Stephen Gonzalez, Rellie Liu, and Jean Chen for their contributions to this project’s administration, organization, and data collection.

## Declaration on the Usage of AI

The authors used OpenAI gpt-4o-mini through Stanford University’s AI Playground platform (https://uit.stanford.edu/service/aiplayground) for assistance in typesetting the manuscript in LaTeX. No other AI tools were used in the research or preparation of the manuscript.

## A Appendix

**Literary Excerpt Stimuli**

**Table A1:** Opening and closing sentences of literary excerpt stimuli.

| Excerpt | Opening and closing sentences |
|---|---|
| 501: Uncle Rock by Dagoberto Gilb | *“In the morning, at his favorite restaurant, Erick got to order his favorite American food, sausage and eggs and hash-brown papitas fried crunchy on top.*<br>*[…]*<br>*He was a salesman in an appliance store who’d bragged that a rich customer had given it to him and so why shouldn’t he give it to Erick’s mom, who couldn’t afford such a good TV otherwise?”* |
| 502: Nothing Living Lives Alone by Wendell Berry | *“Andy Catlett was a child of two worlds.*<br>*[…]*<br>*Though he could not always resist the temptation to be lazy, he knew that laziness was what it was, and he was embarrassed by it even as he indulged in it.”* |
| 503: Corrie by Alice Munro | *“After Corrie’s father died, the shoe factory had been taken over by a large firm that had promised — so she believed — to keep it running.*<br>*[…]*<br>*So she hired him to do the lawns at her house — something she’d been doing herself for the exercise, but her figure didn’t really need it and it took forever with her lameness.”* |
| 504: Mountain Ways by Ursula K. Le Guin | *“In the stony uplands of the Deka Mountains the farmholds are few and far between.*<br>*[…]*<br>*She was a farm woman who had gone to school.”* |
| 505: How the Potato Changed the World by Charles C. Mann (Smithsonian Magazine) | *“Geographically, the Andes are an unlikely birthplace for a major staple crop.*<br>*[…]*<br>*Night winds carry the smell of roasting potatoes for what seems like miles.”* |
| 506: The Forest Unseen by David Haskell | *“The leaf litter is mottled with browns.*<br>*[…]*<br>*Like a farmer tending her apple trees and her field of corn, a lichen is a melding of lives.”* |

## Notes

### Competing Interest Statement

The authors have declared no competing interest.

## References

Bates, D., Mächler, M., Bolker, B., and Walker, S. (2015). Fitting linear mixed-effects models using lme4. Journal of Statistical Software, 67(1):1–48.

Benjamini, Y., Drai, D., Elmer, G., Kafkafi, N., and Golani, I. (2001). Controlling the false discovery rate in behavior genetics research. Behavioural Brain Research, 125(1–2):279–284.

Black, J. E. and Barnes, J. L. (2015). The effects of reading material on social and non-social cognition. Poetics, 52:32–43.

Boyd, R. L., Ashokkumar, A., Seraj, S., and Pennebaker, J. W. (2022). The development and psychometric properties of LIWC-22. Austin, TX: University of Texas at Austin.

Brainard, D. H. (1997). The psychophysics toolbox. Spatial Vision, 10(4):433–436.

Broderick, M. P., Anderson, A. J., Di Liberto, G. M., Crosse, M. J., and Lalor, E. C. (2018). Electrophysiological correlates of semantic dissimilarity reflect the comprehension of natural, narrative speech. Current Biology, 28(5):803–809.

Busselle, R. and Bilandzic, H. (2008). Fictionality and perceived realism in experiencing stories: A model of narrative comprehension and engagement. Communication Theory, 18(2):255–280.

Cheng, A. (2008). Individualized engagement with genre in academic literacy tasks. English for Specific Purposes, 27(4):387–411.

Cohen, S. S. (2018). The Inter-Subject Correlation of EEG in Response to Naturalistic Stimuli. PhD thesis, The City University of New York.

Cohen, S. S., Henin, S., and Parra, L. C. (2017). Engaging narratives evoke similar neural activity and lead to similar time perception. Scientific Reports, 7(1):4578.

Cohen, S. S. and Parra, L. C. (2016). Memorable audiovisual narratives synchronize sensory and supramodal neural responses. ENeuro, 3(6).

Colman, P. (2007). A new way to look at literature: A visual model for analyzing fiction and nonfiction texts. Language Arts, 84(3):257–268.

Dauer, T., Nguyen, D. T., Gang, N., Dmochowski, J. P., Berger, J., and Kaneshiro, B. (2021). Inter-subject correlation while listening to minimalist music: A study of electrophysiological and behavioral responses to Steve Reich’s Piano Phase. Frontiers in Neuroscience, 15:702067.

Davidesco, I., Laurent, E., Valk, H., West, T., Milne, C., Poeppel, D., and Dikker, S. (2023). The temporal dynamics of brain-to-brain synchrony between students and teachers predict learning outcomes. Psychological Science, 34(5):633–643.

Delorme, A. and Makeig, S. (2004). EEGLAB: An open source toolbox for analysis of single-trial EEG dynamics including independent component analysis. Journal of Neuroscience Methods, 134(1):9–21.

Dmochowski, J. P., Greaves, A. S., and Norcia, A. M. (2015). Maximally reliable spatial filtering of steady state visual evoked potentials. NeuroImage, 109:63–72.

Dmochowski, J. P., Sajda, P., Dias, J., and Parra, L. C. (2012). Correlated components of ongoing EEG point to emotionally laden attention–a possible marker of engagement? Frontiers in Human Neuroscience, 6:112.

Dodell-Feder, D., Koster-Hale, J., Bedny, M., and Saxe, R. (2011). fMRI item analysis in a theory of mind task. NeuroImage, 55(2):705–712.

Finn, E. S., Corlett, P. R., Chen, G., Bandettini, P. A., and Constable, R. T. (2018). Trait paranoia shapes inter-subject synchrony in brain activity during an ambiguous social narrative. Nature Communications, 9(1):Article 1.

Grill-Spector, K., Henson, R., and Martin, A. (2006). Repetition and the brain: Neural models of stimulus-specific effects. Trends in Cognitive Sciences, 10(1):14–23.

Hasson, U., Nir, Y., Levy, I., Fuhrmann, G., and Malach, R. (2004). Intersubject synchronization of cortical activity during natural vision. Science, 303(5664):1634–1640.

Holm, S. (1979). A simple sequentially rejective multiple test procedure. Scandinavian Journal of Statistics, pages 65–70.

Honey, C. J., Thesen, T., Donner, T. H., Silbert, L. J., Carlson, C. E., Devinsky, O., Doyle, W. K., Rubin, N., Heeger, D. J., and Hasson, U. (2012). Slow cortical dynamics and the accumulation of information over long timescales. Neuron, 76(2):423–434.

Iotzov, I., Fidali, B. C., Petroni, A., Conte, M. M., Schiff, N. D., and Parra, L. C. (2017). Divergent neural responses to narrative speech in disorders of consciousness. Annals of Clinical and Translational Neurology, 4(11):784–792.

Irsik, V. C., Johnsrude, I. S., and Herrmann, B. (2022). Neural activity during story listening is synchronized across individuals despite acoustic masking. Journal of Cognitive Neuroscience, 34(6):933–950.

Kaneshiro, B., Nguyen, D. T., Norcia, A. M., Dmochowski, J. P., and Berger, J. (2020). Natural music evokes correlated EEG responses reflecting temporal structure and beat. NeuroImage, 214:116559.

Kaneshiro, B., Nguyen, D. T., Norcia, A. M., Dmochowski, J. P., and Berger, J. (2024). Inter-subject correlation of electroencephalographic and behavioural responses reflects time-varying engagement with natural music. European Journal of Neuroscience, 59(12):3162–3183.

Kaufeld, G., Bosker, H. R., ten Oever, S., Alday, P. M., Meyer, A. S., and Martin, A. E. (2020). Linguistic structure and meaning organize neural oscillations into a content-specific hierarchy. Journal of Neuroscience, 40(49):9467–9475.

Ki, J. J., Kelly, S. P., and Parra, L. C. (2016). Attention strongly modulates reliability of neural responses to naturalistic narrative stimuli. Journal of Neuroscience, 36(10):3092–3101.

Kidd, D. C. and Castano, E. (2013). Reading literary fiction improves theory of mind. Science, 342(6156):377– 380.

Kidd, D. C. and Castano, E. (2019). Reading literary fiction and theory of mind: Three preregistered replications and extensions of kidd and castano (2013). Social Psychological and Personality Science, 10(4):522–531.

Kleiner, M., Brainard, D., and Pelli, D. (2007). What’s new in Psychtoolbox-3? https://pure.mpg.de/rest/items/item_1790332/component/file_3136265/content. Accessed 2026-09-30.

Lahnakoski, J. M., Glerean, E., Jääskeläinen, I. P., Hyönä, J., Hari, R., Sams, M., and Nummenmaa, L. (2014). Synchronous brain activity across individuals underlies shared psychological perspectives. NeuroImage, 100:316–324.

Lerner, Y., Honey, C. J., Silbert, L. J., and Hasson, U. (2011). Topographic mapping of a hierarchy of temporal receptive windows using a narrated story. Journal of Neuroscience, 31(8):2906–2915.

Malave, A. J., Hernandez, P. A., and Kaneshiro, B. (2026). SENSI-EEG-Preproc-CS: A MATLAB preprocessing pipeline for EEG responses to continuous stimuli. Stanford Digital Repository.

Mar, R. A. and Oatley, K. (2008). The function of fiction is the abstraction and simulation of social experience. Perspectives on Psychological Science, 3(3):173–192.

Nastase, S. A., Gazzola, V., Hasson, U., and Keysers, C. (2019). Measuring shared responses across subjects using intersubject correlation. Social Cognitive and Affective Neuroscience, 14(6):667–685.

Nguyen, M., Vanderwal, T., and Hasson, U. (2019). Shared understanding of narratives is correlated with shared neural responses. NeuroImage, 184:161–170.

Norman, G. (2010). Likert scales, levels of measurement and the “laws” of statistics. Advances in Health Sciences Education, 15:625–632.

O’Connell, R. G., Dockree, P. M., Robertson, I. H., Bellgrove, M. A., Foxe, J. J., and Kelly, S. P. (2009). Uncovering the neural signature of lapsing attention: Electrophysiological signals predict errors up to 20 s before they occur. Journal of Neuroscience, 29(26):8604–8611.

Parra, L. C., Spence, C. D., Gerson, A. D., and Sajda, P. (2005). Recipes for the linear analysis of EEG. NeuroImage, 28(2):326–341.

Petroni, A., Cohen, S. S., Ai, L., Langer, N., Henin, S., Vanderwal, T., Milham, M. P., and Parra, L. C. (2018). The variability of neural responses to naturalistic videos change with age and sex. ENeuro, 5(1):ENEURO.0244–17.2017.

Posit Team (2025). RStudio: Integrated development environment for R. Posit Software, PBC.

Poulsen, A. T., Kamronn, S., Dmochowski, J., Parra, L. C., and Hansen, L. K. (2017). EEG in the classroom: Synchronised neural recordings during video presentation. Scientific Reports, 7(1):Article 1.

Rosenkranz, M., Holtze, B., Jaeger, M., and Debener, S. (2021). EEG-based intersubject correlations reflect selective attention in a competing speaker scenario. Frontiers in Neuroscience, 15:685774.

Saxe, R., Carey, S., and Kanwisher, N. (2004). Understanding other minds: Linking developmental psychology and functional neuroimaging. Annual Review of Psychology, 55(1):87–124.

Silbert, L. J., Honey, C. J., Simony, E., Poeppel, D., and Hasson, U. (2014). Coupled neural systems underlie the production and comprehension of naturalistic narrative speech. Proceedings of the National Academy of Sciences, 111(43):E4687–E4696.

Simony, E., Honey, C. J., Chen, J., Lositsky, O., Yeshurun, Y., Wiesel, A., and Hasson, U. (2016). Dynamic reconfiguration of the default mode network during narrative comprehension. Nature Communications, 7(1):12141.

Stephens, G. J., Silbert, L. J., and Hasson, U. (2010). Speaker–listener neural coupling underlies successful communication. Proceedings of the National Academy of Sciences, 107(32):14425–14430.

Tompson, S. H., Kahn, A. E., Falk, E. B., Vettel, J. M., and Bassett, D. S. (2019). Individual differences in learning social and nonsocial network structures. Journal of Experimental Psychology: Learning, Memory, and Cognition, 45(2):253–271.

Tucker, D. M. (1993). Spatial sampling of head electrical fields: The geodesic sensor net. Electroencephalography and Clinical Neurophysiology, 87(3):154–163.

Ueno, F. and Shimada, S. (2023). Inter-subject correlations of EEG reflect subjective arousal and acoustic features of music. Frontiers in Human Neuroscience, 17.

Van den Broek, P. and Helder, A. (2017). Cognitive processes in discourse comprehension: Passive processes, reader-initiated processes, and evolving mental representations. Discourse Processes, 54(5–6):360–372.

Wilson, S. M., Molnar-Szakacs, I., and Iacoboni, M. (2008). Beyond superior temporal cortex: Intersubject correlations in narrative speech comprehension. Cerebral Cortex, 18(1):230–242.

Yeshurun, Y., Swanson, S., Simony, E., Chen, J., Lazaridi, C., Honey, C. J., and Hasson, U. (2017). Same story, different story: The neural representation of interpretive frameworks. Psychological Science, 28(3):307–319.

